# Data-driven speciation tree prior for better species divergence times in calibration-poor molecular phylogenies

**DOI:** 10.1101/2021.03.27.437326

**Authors:** Qiqing Tao, Jose Barba-Montoya, Sudhir Kumar

## Abstract

**Motivation:** Precise time calibrations needed to estimate ages of species divergence are not always available due to fossil records’ incompleteness. Consequently, clock calibrations available for Bayesian dating analyses can be few and diffused, i.e., phylogenies are calibration-poor, impeding reliable inference of the timetree of life. We examined the role of speciation birth-death tree prior on Bayesian node age estimates in calibration-poor phylogenies and tested the usefulness of an informative, data-driven tree prior to enhancing the accuracy and precision of estimated times.

**Results:** We present a simple method to estimate parameters of the birth-death tree prior from the molecular phylogeny for use in Bayesian dating analyses. The use of a data-driven birth-death (ddBD) tree prior leads to improvement in Bayesian node age estimates for calibration-poor phylogenies. We show that the ddBD tree prior, along with only a few well-constrained calibrations, can produce excellent node ages and credibility intervals, whereas the use of an uninformative, uniform (flat) tree prior may require more calibrations. Relaxed clock dating with ddBD tree prior also produced better results than a flat tree prior when using diffused node calibrations. We also suggest using ddBD tree priors to improve the detection of outliers and influential calibrations in cross-validation analyses.

**Conclusion:** Empirical Bayesian dating analyses with ddBD tree priors enable more accurate and precise node age estimates for calibration-poor phylogenies. Our results have practical applications because the ddBD tree prior reduces the number of well-constrained calibrations necessary to obtain reliable node age estimates. This would help address key impediments in building the grand timetree of life, revealing the process of speciation, and elucidating the dynamics of biological diversification.

**Availability:** An R module for computing the ddBD tree prior, simulated datasets, and empirical datasets are available at https://github.com/cathyqqtao/ddBD-tree-prior.

## 1. Introduction

In Bayesian relaxed clock methods for estimating species divergence times from molecular sequences, node age estimates are a product of the interaction of calibration time priors and the speciation tree prior applied to a molecular phylogeny (Barba-Montoya *et al*., 2017; Warnock *et al*., 2012; Yang, 2006). The calibration time priors come from incomplete fossil-record, often resulting in phylogenies with only a few clock calibrations that are well-constrained (Hipsley and Müller, 2014; Parham *et al*., 2012). Even in fossil-rich taxonomic groups, the quality and quantity of calibration vary across clades, and node calibrations are missing in most clades. Actually, most phylogenies have only a few (or even no) well-constrained calibrations. So, molecular clock dating studies resort to using calibrations derived from previous molecular studies or use pre-determined evolutionary rates to convert sequence divergences to time due to a paucity of calibrations (Hipsley and Müller, 2014; Tao *et al*., 2020).

Obviously, an increase in the quantity and quality of clock calibrations will remedy many problems facing molecular dating today. However, it is not easy or cost-effective to acquire additional fossil records and other information sources to establish node calibrations. An alternative candidate to improve divergence time estimates in calibration-poor phylogenies is the use of informative tree priors for speciation, which is theoretically expected to improve the Bayesian estimates of node ages. For example, in the MCMCTree software (Yang, 2007), a birth-death (BD) speciation tree prior consists of a per-lineage birth rate (λ) and death rate (μ) of the birth-death process as well as a species sampling fraction (ρ). Parameters λ and μ are the numbers of new species arising and going extinct in a given time unit, respectively, and ρ is the fraction of sampled species in the phylogeny.

For a given phylogeny of species (only the relationships), the given set of parameters of speciation tree prior predicts a distribution of node times (**Fig. 1**). **Figure 1A** shows two distinct tree priors produced by different combinations of parameter values. The flat tree prior, which investigators often use, has a uniform-like density. It posits that deep and recent node ages have a similar probability of occurring anywhere in the phylogeny (**Fig. 1B**). In contrast, a skewed tree prior may posit a higher probability for having recent divergences and a lower probability of deep divergences in the phylogeny (e.g., **Fig. 1C**) or vice versa.

**Figure 1.**
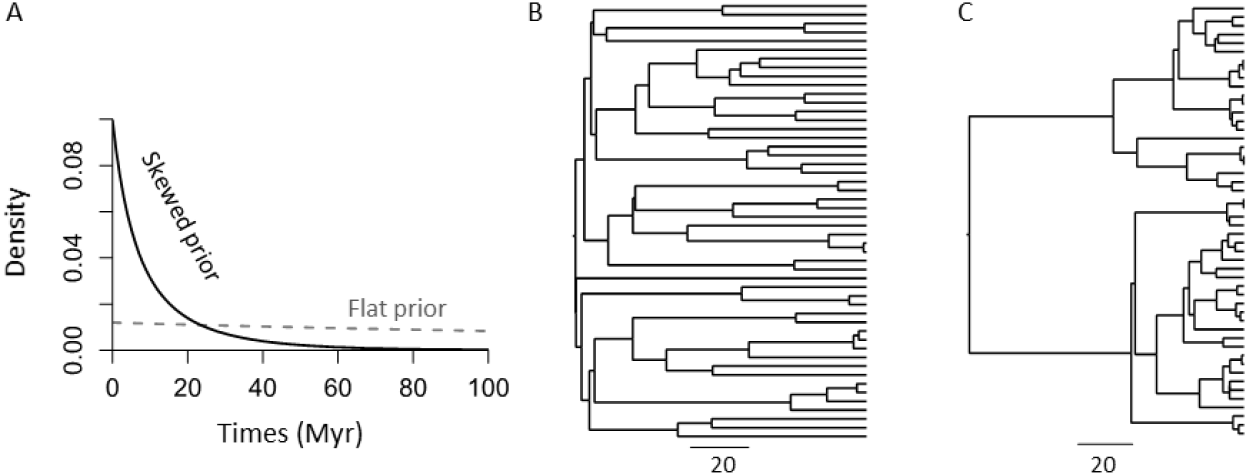
(A) Distinct density plots for a flat and a skewed tree prior. An example timetree simulated under the birth-death process using (B) a flat and (C) a skewed tree prior is shown. Both timetrees have root age of 100 million years. Birth rate, death rate, and sampling fraction are (0.02, 0.02, 0.1) and (0.1, 0.05, 0.99) in the simulation for flat prior and skewed prior, respectively.

It is well-appreciated that the choice of the tree prior has a limited impact on time estimates when many well-constrained calibrations are applied because the calibration time prior exerts a strong influence on estimated times (Barba-Montoya *et al*., 2018; Foster *et al*., 2016). However, the choice of the speciation tree prior is expected to influence node age estimates in the absence of informative calibrations (Yang and Rannala, 1997; Heled and Drummond, 2012, 2015; dos Reis *et al*., 2018; Yang, 2006). Interestingly, most investigators also use a flat tree prior for calibration-poor phylogenies, which is the default setting in MCMCTree and other Bayesian software (Ronquist and Huelsenbeck, 2003). However, if the actual distribution of node times is not flat (uniform-like), posterior time estimates may be biased in calibration-poor phylogenies, as there is not enough information to constrain posterior estimates.

Therefore, we wondered whether using an informative tree prior can rescue time estimates in calibration-poor phylogenies. A survey of the literature revealed no past investigations that examined the power of an informative tree prior to dating calibration-poor phylogenies. We only found studies to have investigated the impact of the tree prior for analyzing empirical datasets, rather than simulated datasets, and with many well-constrained calibrations (Barba-Montoya *et al*., 2018). Neither of those situations offers insight into informative speciation tree prior’s usefulness in dating calibration-poor phylogenies.

So, we assessed the benefit of using the known (correct) tree prior by analyzing a simulated dataset. We found that the node ages were overestimated by 222% when the default tree prior was used, i.e., estimates were highly biased (both average absolute error and average error are 222%). The estimated highest posterior density intervals (HPDs) only contained the true times for 14.6% of nodes, i.e., a coverage probability of 14.6%. In sharp contrast, the use of correct tree prior reduced the average absolute error of node ages to 33.7% and average error to 30.0% and increased the HPD coverage probability to 95.8%. This simple example suggests that using tree prior may improve posterior point time estimates and HPDs (**Fig. 2**; see *Results* for more details and expanded discussion).

**Figure 2.**
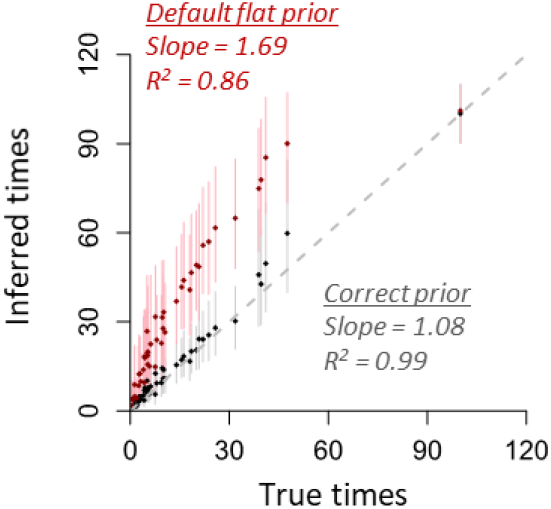
Comparison of true times with node ages inferred using the correct (black) and the default flat (red) tree prior for the dataset simulated using model tree in figure 1C. The slope and coefficient of determination (*R*^2^) for the linear regression through the origin are shown. Dots are point estimates of node ages and lines are highest posterior density intervals (HPDs). Gray dashed line is 1:1 line.

This prompted us to devise a simple method to estimate tree prior’s parameters based solely on a molecular phylogeny with branch lengths (number of substitutions per site). In the following, we first present this approach to generate data-driven estimates of BD (ddBD) tree prior to use in Bayesian relaxed clock methods implemented in MCMCTree. This method is an empirical Bayesian approach. We then explore ddBD tree priors’ use in analyzing empirical and simulated datasets to quantify increased accuracy, decreased bias, and higher coverage probabilities of Bayesian estimate in calibration-poor phylogenies. We consider well-constrained calibrations and diffused uncertainty densities of calibrations to test if ddBD tree priors produce better estimates than the default (flat) tree prior in MCMCTree. We investigate the impact of using an incorrect substitution model and phylogenies on the improvement in divergence times offered by the use of ddBD tree priors. We also discuss the usefulness of ddBD tree priors in additional practical situations.

## 2. Method and Materials

### 2.1. A simple approach for estimating parameters of birth-death speciation tree prior

A simple approach for building the tree prior by using a species phylogeny and multispecies sequence alignment is to estimate node ages using the relative rate framework (RRF) without calibrations first (Tamura *et al*., 2012, 2018). RRF is a relaxed-clock method that does not require the specification of any priors, such as calibration, clock rate, and tree priors. RRF is selected because it has been shown to produce reliable relative node age estimates under various scenarios and conditions (Tamura *et al*., 2012, 2018). However, other non-Bayesian methods, e.g., penalized likelihood method (Sanderson, 2002), can also be used to estimate node ages.

We utilize the kernel density distribution of node times generated by RRF to estimate λ, μ, and ρ needed by the MCMCTree program using the following equations (Yang, 2006) through a maximum likelihood procedure.

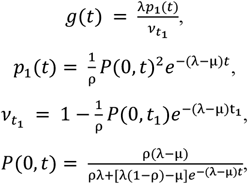

where *g*(*t*) is the kernel density of node times, *p*_1_(*t*) is the probability that a lineage arising in the past leaves exactly one descendant in the sample, *P*(0,*t*) is the probability that a lineage arising at time *t* in the past leaves one or more descendants in the present-day sample, and *t*_1_ is the root age. In our calculation, we first assume the root age is 1, so we estimate relative values of λ, μ, and ρ. Then, we scale the relative parameter values based on the root node’s age, which is required by the MCMCTree program. As there are no analytical solutions to estimate all three parameters simultaneously (Stadler, 2009), so we estimate them numerically.

Because maximum likelihood estimation’s performance relies on good initial values, we conduct a three-dimensional grid search of the initial values. The grid search involves ten λ values (1.1 – 10.1, step size = 1), ten μ values (1 – 10, step size = 1), and five ρ values to represent extremely sparse to highly dense sampling (0.001, 0.01, 0.1, 0.5, and 0.9). We set λ > μ, a biologically reasonable constraint for a phylogeny of living species. These parameter choices in grid search ensure that a wide range of tree prior densities is explored. From these 500 parameter choices, we select the set that maximizes the predicted density’s fit with the distribution of RRF node times as the initial value. The fitness of predicted density can be determined by the sum of squared errors or Kullback-Leibler divergence (Kullback and Leibler, 1951), both of which produced similar results. We used the sum of squared errors in all analyses reported here. The grid search procedure was very fast (<1 second for a tree of 274 taxa in **Fig. 3**). Also, it has the flexibility to specify ρ if the species sampling fraction is known. In this case, only λ and μ need to be estimated. Using the best estimates of λ, μ, and ρ, we conduct a maximum likelihood analysis to obtain the final estimates of λ, μ, and ρ.

**Figure 3.**
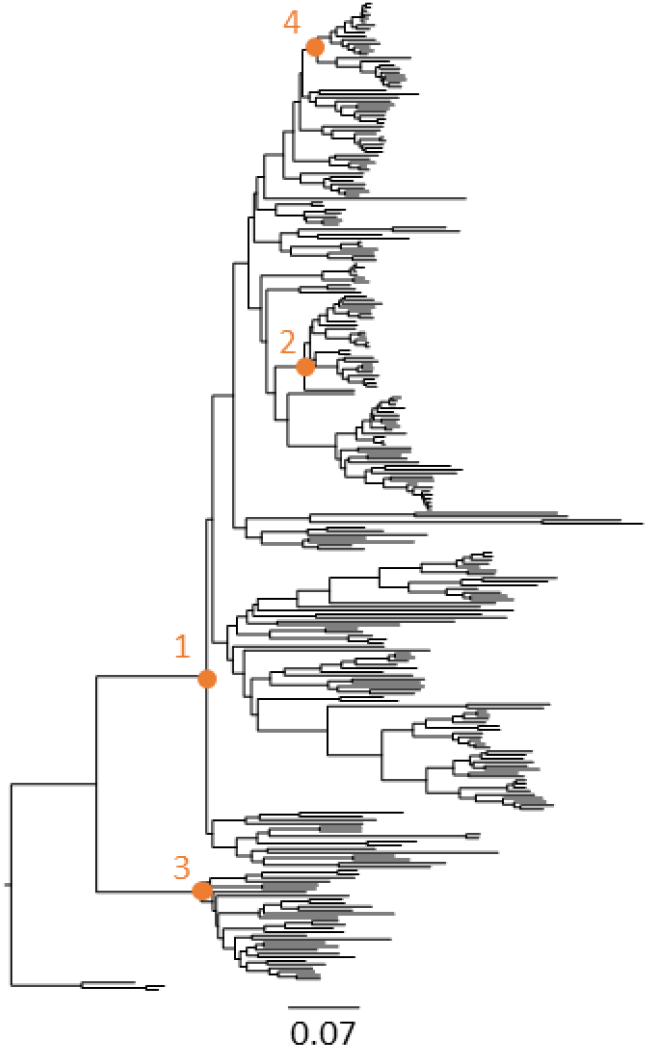
Evolutionary relationships and branch lengths derived from an alignment of 7,370 base pairs of 12 mitochondrial genes from 274 mammalian species from dos Reis *et al*. (2012). Orange nodes were calibrated in the evaluations in which fewer calibrations were utilized. The number indicates the order of calibration addition.

We must note that many combinations of the three parameters can produce the same tree prior density, so the estimates produced are not unique, but this will not impact MCMCTree time estimates (see *Results* and *Discussion*).

### 2.2. Simulated datasets and analysis

We simulated ten model timetrees of 50 tips under the birth-death process with λ, μ, and ρ of 10, 5, and 0.99 (time unit = 100 million years, Myr) using “sim.bd.taxa.age” function in TreeSim R package (Stadler, 2011). The age of the most recent common ancestor (MRCA) of species was fixed to be 100 Myr. An example model tree is shown in **Fig. 1C**. We then simulated molecular evolutionary rates to generate a phylogram for each model timetree. An example phylogram is shown in **Fig. 4A**. Branch-wise evolutionary rates were drawn from a lognormal distribution where the mean rate was 0.00228 substitutions per site per Myr, and the variance was 0.4 (log-scale) (**Fig. 4B**). Rates varied independently among sites.

**Figure 4.**
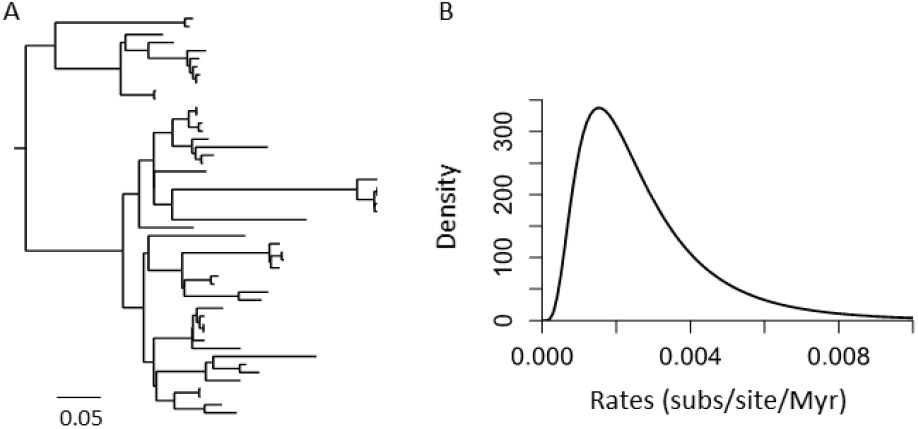
(A) An example simulated branch length tree. (B) Distribution used in the computer simulation to sample molecular evolutionary rates for branches.

Finally, alignments of 8,000 base pairs (bp) were generated based on phylograms in SeqGen (Rambaut and Grassly, 1997) under the Hasegawa-Kishino-Yano (HKY) (Hasegawa *et al*., 1985) substitution model with the assumption of substitution rate variation across sites under a gamma distribution (+G, α = 0.4). All parameter values used in simulations were derived from an empirical mammalian study (dos Reis *et al*., 2012). We need time estimates of all branches, but RelTime does not produce estimates for the rooting outgroup. So, we added an arbitrary rooting outgroup to all model timetrees for simulating sequences. The arbitrary sequence of rooting outgroup was later removed in MCMCTree analyses.

We estimated divergence times using the default flat (uniform-like) BD setting (2 2 0.1) and the ddBD tree prior for the correct tree topology. Three calibration strategies were used: (a) root calibration only, (b) root and one internal node calibration (root + 1C), and (c) root and three internal node calibrations (root + 3C). In root calibration only analysis, we used a uniform density (true age ± 10 Myr) to calibrate root age. Besides the default flat and ddBD tree priors, we also analyzed datasets with only the root calibration with the simulated tree prior. In the root+1C analysis, we randomly selected one internal node to calibrate, along with the root calibration. This calibrated node may come from shallow, intermediate, and deep regions in the given phylogeny, which enabled us to explore the impact of the calibration location on time estimation. Shallow nodes were nodes with two descending tips; deep nodes were immediate descendants of the root; intermediate nodes are nodes whose ages were 30%-70% of the root age. A uniform density (true age ± 50% of true age) was used to calibrate the selected shallow, intermediate, and deep nodes. In the root+3C calibration analysis, we applied the shallow, intermediate, and deep calibration together with the root calibration in each phylogeny.

We also test our method’s performance when the phylogenetic tree has an error and when an oversimplified model of nucleotide substitution is used to assess how the improvement in time estimates using ddBD is impacted by such errors. We used the Jukes-Cantor (JC) model (Jukes and Cantor, 1969) as the incorrect model, which is the most extreme violation of the HKY+G substitution model when considering the general time-reversible models. Comparison of time estimates using the JC model and HKY+G model with and without an informative prior is interesting to examine whether substitution model violation has a more serious impact on time estimates using the ddBD. In these analyses, we also compared the detrimental impact of using inferred phylogenies rather than correct phylogenies when estimating ddBD. Neighbor-joining (NJ) method (Saitou and Nei, 1987) with JC and Tamura-Nei (TrN) (Tamura and Nei, 1993) model were used to infer phylogenies in MEGA X (Kumar *et al*., 2018). We used the TrN model with a gamma distribution (+G) of rate variation among sites in tree inference, as are no analytical solutions for pairwise distance calculations using the HKY model, and it is nested within the TrN model for which an analytical solution exists. These inferred phylogenies contained many topological differences from the correct tree. We compared estimated times with the true times for eight different combinations (flat vs. ddBD tree prior, simple vs. complex substitution model, and inferred vs. correct phylogeny). Because inferred phylogenies are not identical to the correct trees used in the simulation, time estimates compared were the pairwise species divergence times in the true timetree and the inferred Bayesian inferred timetree.

In all dating analyses, we assigned to the overall rate (μ) a gamma hyperprior G(1, 4) with a mean of 0.25, which translates into a rate of 0.0025 substitutions per site per Myr that is similar to the true value. For the rate drift parameter (σ2), we assigned another gamma hyperprior G(1, 10) with a mean of 0.1. Two independent runs of 2,000,000 generations were conducted, and results were checked for convergence (ESS >200). The performance was assessed by mean absolute percent error (MAPE) over all nodes for each dataset to measure the accuracy of time estimates, mean percent error (MPE) to measure the bias of time estimates over all nodes in phylogeny, and the HPD coverage (i.e., percent of true node times contained in estimated HPDs) to measure the coverage probability of HPDs across all nodes in a phylogeny.

### 2.3. Empirical dataset and analysis

We inferred the tree prior parameters for a mammalian dataset of 274 species (7,370 bp) obtained from dos Reis et al. (2012). The original phylogeny (**Fig. 3**) and the alignment were used to estimate node ages in MCMCTree v4.7h (Yang 2007). We also estimated node ages using the default flat tree prior used in the original study (1 1 0; uniform density). All other priors were specified as in the original study. Thirty-five internal calibrations were used in the original study. We considered 10% of original count of calibrations as the calibration-poor case (i.e., internal calibration count ≤ 4), Therefore, we applied five calibration strategies: (a) root calibration only, (b) root + 1C, (3) root + 2C, (4) root + 3C, and (5) root + 4C internal node calibrations. In the root calibration analysis and other analyses, we used the original study’s root calibration. In the root + 1C, we selected a deep calibration from the 35 calibrations used in the original study because the simulated analysis has shown that the deep calibration has a bigger impact on time estimation. In the root + 2C analysis, we randomly selected an internal calibration and used it together with the one used in the one-calibration analysis. A similar strategy for choosing calibrations was used for root + 3C and root + 4C analyses.

To assess the impact of using diffused calibrations, we conducted analyses using the same calibrated nodes but applying wider boundaries. Calibrations were specified using a uniform distribution U(t_L_, t_U_), where t_L_ is the minimum age bound and t_U_ the maximum age bound. The offset value from the original skewed student t (ST) distribution was used as the minimum bound (t_L_), and the maximum bound was specified at t_U_ = t_L_+ t_L_/2, which assigns a probability of 50% for the minimum bound (t_L_) to be older. All estimated node ages were compared to the times reported in Tao *et al*. (2020), because the new version of MCMCTree produced times slightly different from those reported in the original study (v4.2e was used). For each analysis, two independent runs of at least 1,500,000 generations were conducted, posterior estimates were checked for convergence (ESS values >200).

## 3. Results

### 3.1. Correct tree priors enable better time estimates

We first tested whether a known tree prior is beneficial. We used the simulated tree prior (i.e., correct tree prior) and the default flat tree prior in the MCMCTree analysis of the simulated datasets (see *Method and Materials*). No calibration priors were placed on the phylogeny’s internal nodes to focus purely on the impact of speciation tree priors.

Node ages were estimated with rather large errors when using the flat prior (108% −334%, **Fig. 5A** and **B**), and HPDs only contained the true times for a few nodes (coverage probability = 6.3% −54.2%, **Fig. 5C**). MAPE and MPE of flat tree prior estimates were similar because the use of flat prior overestimated the node times for all the simulated datasets. An uninformative flat tree prior produced highly biased estimates in the absence of calibrations in these analyses. It is because the flat tree prior tends to assign a similar number of shallow and deep node times on a phylogeny that was simulated using a skewed tree prior and contains more shallow nodes (**Fig. 1A**).

**Figure 5.**
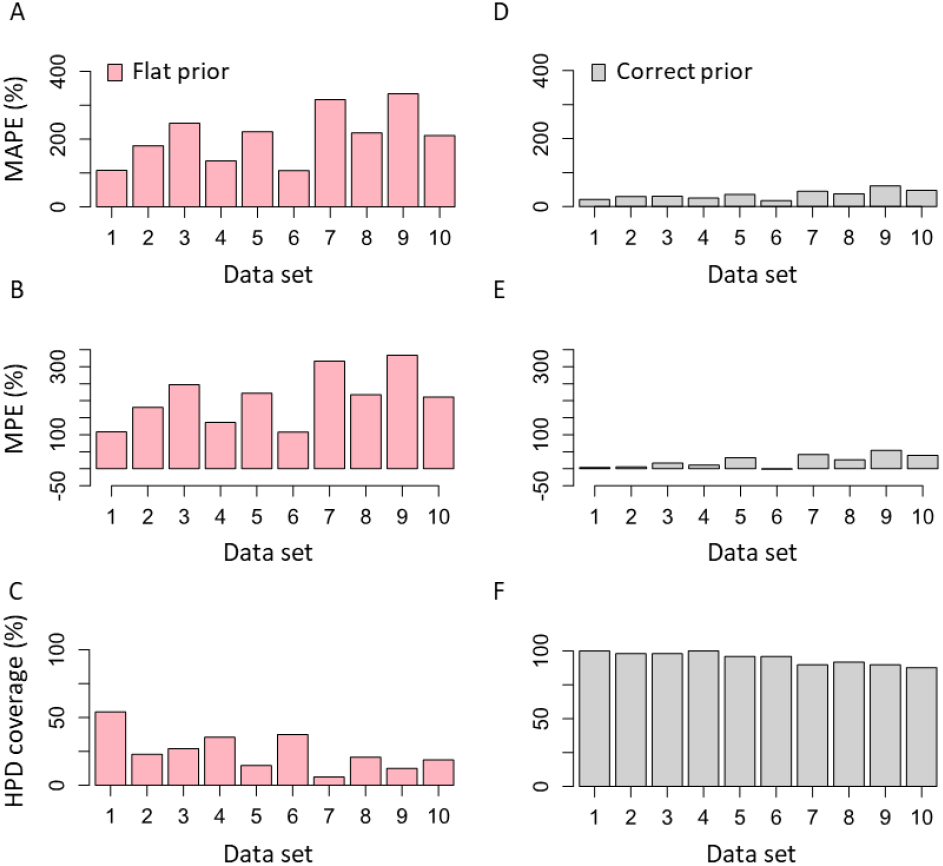
(A, D) Mean of absolute percent error, MAPE, (B, E) mean of percent error, MPE, and (C, F) HPD coverage for nodes times estimated using the default flat (red) and correct (gray) tree prior for each dataset.

The use of correct tree prior was very beneficial, as the mean absolute error (17.2% −61.1%, **Fig. 5D**) and mean error (−1.6% −54.4%, **Fig. 5E**) were reduced tremendously, and HPD coverage became much higher (87.8% −100%**; Fig. 5F**). Estimates with large errors were usually found at recent nodes (< 5 Myr), as the number of substitutions is limited. However, these results clearly show that tree prior selection can significantly impact node age estimates. If a suitable tree prior can be estimated *a priori*, one will likely obtain much better time estimates even without using internal calibrations (**Fig. 2** and **5**).

### 3.2. Accuracy of ages with ddBD tree prior

Analysis of simulated datasets using ddBD tree prior produced time estimates comparable to those obtained using the correct priors for all datasets (**Fig. 6**). Using ddBD priors, we achieved MAPE of node age as low as 17.1% (the flat prior MAPE was 107.7%), whereas the maximum MAPE was 64.2% compared to 333.9% for the flat prior. Clearly, ddBD performed better for some datasets than others, but this was not due to difficulties in estimating tree prior parameters, as the use of correct priors produced a similar amount of error.

**Figure 6.**
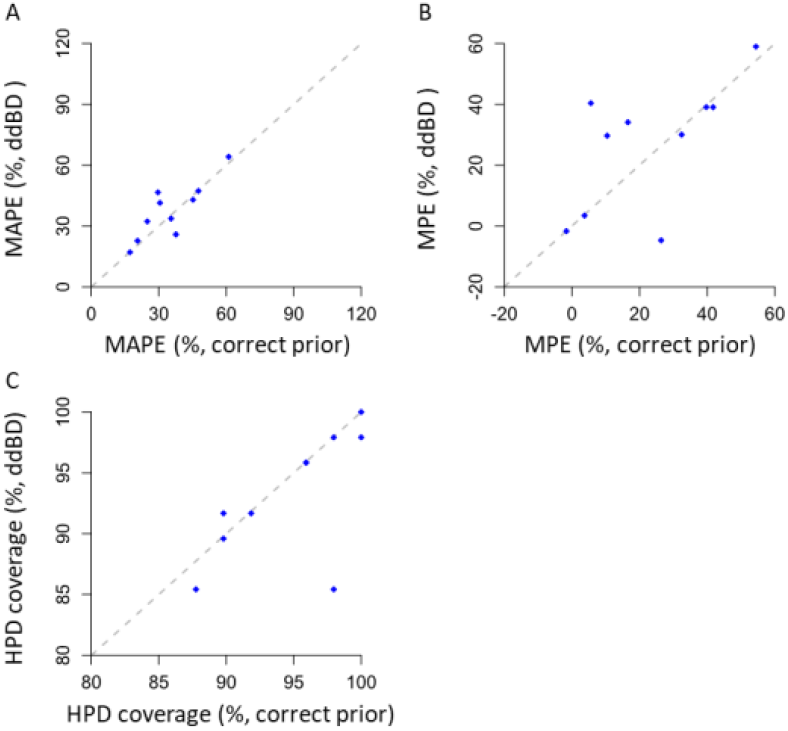
Comparison of (A) mean of absolute percent error, MAPE, (B) mean of percent error, MPE, and (C) HPD coverage for time estimates obtained using correct and ddBD tree prior for each dataset. Only the root calibration was used. Dashed line is 1:1 line.

The comparison of mean percent error (MPE) of node ages showed a similar trend, where the MPE of node times estimated using ddBD tree prior (−1.7% −59.0%) was significantly smaller than the estimates from the flat prior (107.7% −333.9%). The improvement in HPD coverage probability was much more uniform across datasets (>85%), suggesting that it is better to use HPDs estimated using ddBD tree prior in making biological conclusions for calibration-poor phylogenies. These results indicate that the use of ddBD generates less biased estimates.

To assess the generality of the pattern observed in the analysis of simulated datasets, we used ddBD to analyze an empirical alignment (7,370 base pairs) of 12 mitochondrial genes from 274 mammalian species (dos Reis *et al*., 2012) (**Fig. 3**). Since there are no true times in empirical analysis, we compared estimates obtained using ddBD and flat tree priors with those reported in the original study in which 36 calibrations and a flat tree prior were used. In both analyses, we used the sequence alignment, phylogenetic relationships, substitution model (HKY+G_4_), and the root time calibration applied in the original study to enable a direct comparison of absolute times and avoid any confounding effect.

Our approach to estimate ddBD parameters showed an L-shape density of node times (**Fig. 7**, blue curve). This is significantly different from the flat tree prior applied in the original study (*P* < 0.01; **Fig. 7**, red curve). Compared with reported times obtained using 35 well-constrained calibrations, we found that node ages generated using ddBD tree prior and only a root calibration (i.e., no internal calibration) were much more similar (slope = 1.31, **Fig. 7B**) than those produced by using a flat tree prior (slope = 1.94, **Fig. 7B**). It suggested that the use of ddBD can reduce about 60% of differences caused by using a flat tree prior when no internal calibrations are used, which is also congruent with our results of simulated datasets. Therefore, the Bayesian method’s performance with our ddBD tree prior and no internal calibrations is encouraging.

**Figure 7.**
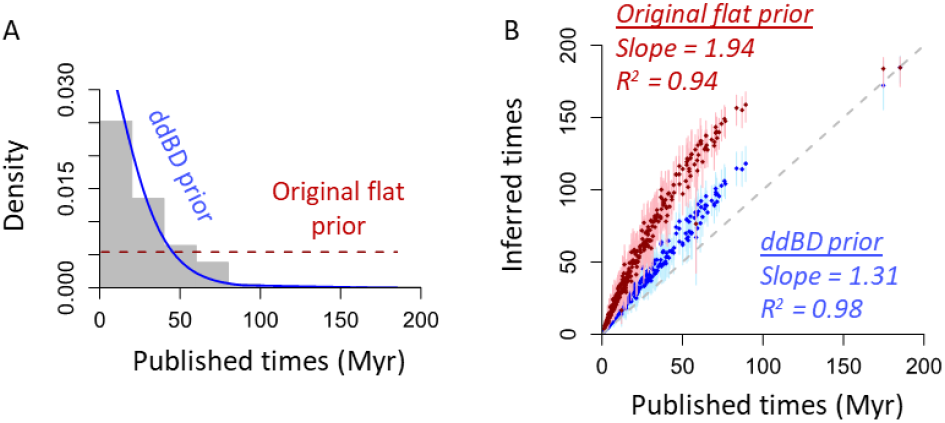
Effect of assumed tree priors on time estimates for the phylogeny in figure 3. (A) Comparison of the ddBD (blue) and original flat (red) tree prior. Gray bars show the distribution of published times. (B) Comparison of node ages inferred using the ddBD (blue) and flat (red) tree prior. The slope and coefficient of determination (*R*^2^) for the linear regression through the origin are shown for simplicity (a second degree polynomial fits the curve better). Dots are point estimates of node ages and lines are HPDs.

### 3.3. Using ddBD tree prior with well-constrained calibrations

The use of ddBD tree prior greatly reduces the difference in node age estimates obtained without internal calibrations compared to those obtained with 35 calibrations for the mammalian dataset. Interestingly, two estimates showed a curvilinear but monotonic relationship (**Fig. 7B**). Therefore, we explored whether the use of only one well-constrained internal calibration can reduce this difference (root+1C). Indeed, concordance increased significantly, as the average difference reduced to less than half (19.0% compared to 46.1%, **Fig. 8A**) and coverage probability doubled (39.0% to 76.1%; **Fig. 8B**). The concordance also increased for estimates obtained using the flat tree prior, but its performance was still worse than the ddBD tree prior (**Fig. 8**).

**Figure 8.**
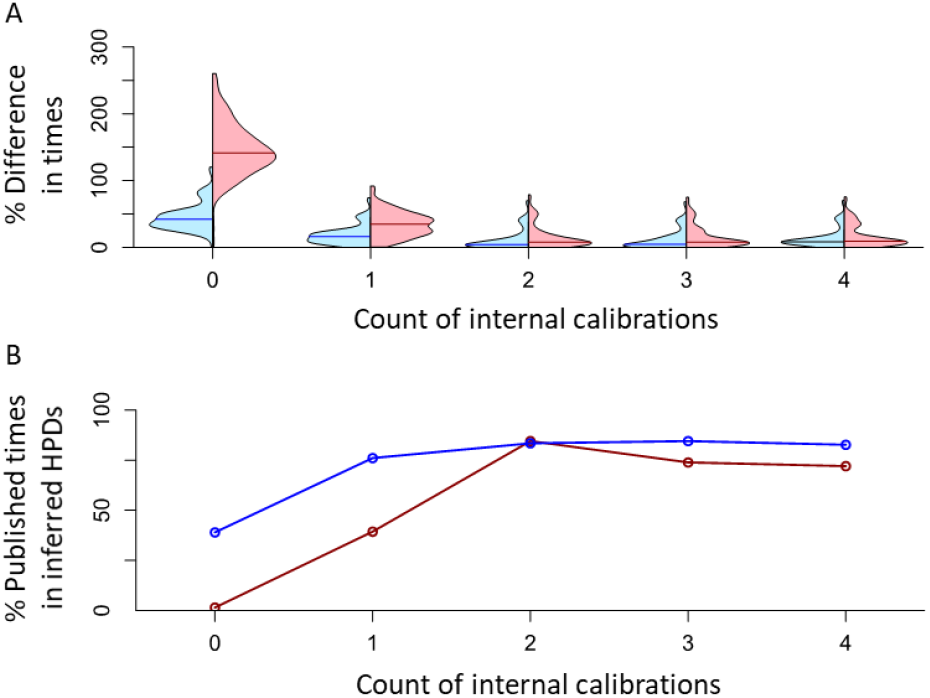
Analysis of empirical data using different numbers of well-constrained internal calibrations. (A) Distribution of absolute percent differences between published times and times estimated using ddBD (blue) and the flat (red) tree prior when different numbers of internal calibrations are used. Lines in violin plots are median values. (B) Percent of nodes whose published times within HPDs obtained using ddBD (blue) and the flat (red) tree prior when different numbers of internal calibrations are used. Zero on x-axis represents the case where only the root calibration and no internal calibrations are used.

These observations prompted us to explore the performance gains made by adding another calibration (root+2C), which made the results very close to those obtained using 35 calibrations in the original study. The difference in time estimates reduces to 11.4%, and the coverage probabilities become higher (83.4%, **Fig. 8**). The addition of a few more well-constrained calibrations only improved the results slightly for ddBD estimates. The ddBD tree’s performance prior was usually better than the performance of the flat tree prior, except for the root+2C case where flat and ddBD tree priors performed similarly.

Overall, the use of fewer than 10% of the internal calibrations (i.e., < 4) along with the ddBD tree prior produced time estimates that differed less than 12% from those reported using all the calibrations. Interestingly, HPD widths for dates with 0-4 internal calibrations were very similar (not shown), which means that estimates and their uncertainties can be generated using a small number of calibrations and ddBD tree prior. We find these results highly encouraging for improving molecular dating in situations where only one or few well-constrained calibrations are known. We confirmed the patterns observed for the empirical data by analyzing simulated datasets (**Fig. 9**). The average error decreased when using one internal calibration and ddBD tree prior (**Fig. 9A**). The median value of the average absolute errors reduced from 37.6% to 29.6%. The HPDs also became slightly better with higher coverage probabilities (improving from 93.1% to 95.8% on average, **Fig. 9B**) with similar HPD widths.

**Figure 9.**
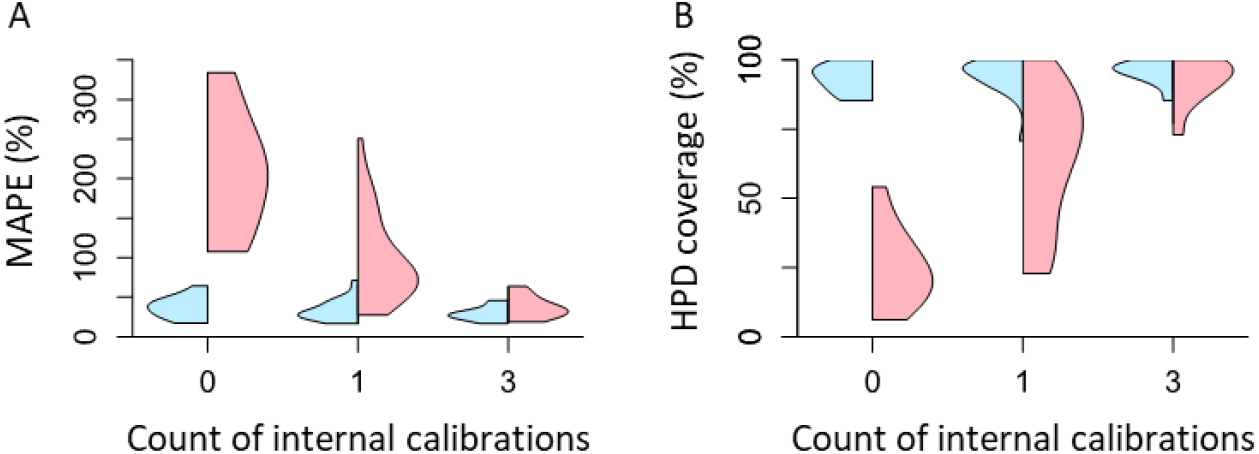
Analysis of simulated datasets using different numbers of well-constrained internal calibrations. Changing pattern of (A) mean absolute percent error, MAPE and (B) mean HPD coverage for estimates obtained using ddBD (blue) and the flat (red) tree prior across all simulated datasets when different numbers of internal calibrations are used. Zero on x-axis represents the case where only the root calibration (i.e., no internal calibrations) are used.

We found that the actual increase in performance depended on the location of the node calibrated, with the highest performance gain seen when deeper nodes were calibrated (**Fig. 10**). This observation is consistent with previous studies where shallow calibrations led to more biased time estimates (van Tuinen and Torres, 2015; Tao *et al*., 2020). Interestingly, when more calibrations were used, the performance of the ddBD tree prior only improved marginally (**Fig. 9**). In contrast, results from the use of flat tree prior always performed worse than ddBD tree prior in all calibration-poor scenarios (**Fig. 9**), consistent with the pattern seen in the empirical data analysis.

**Figure 10.**
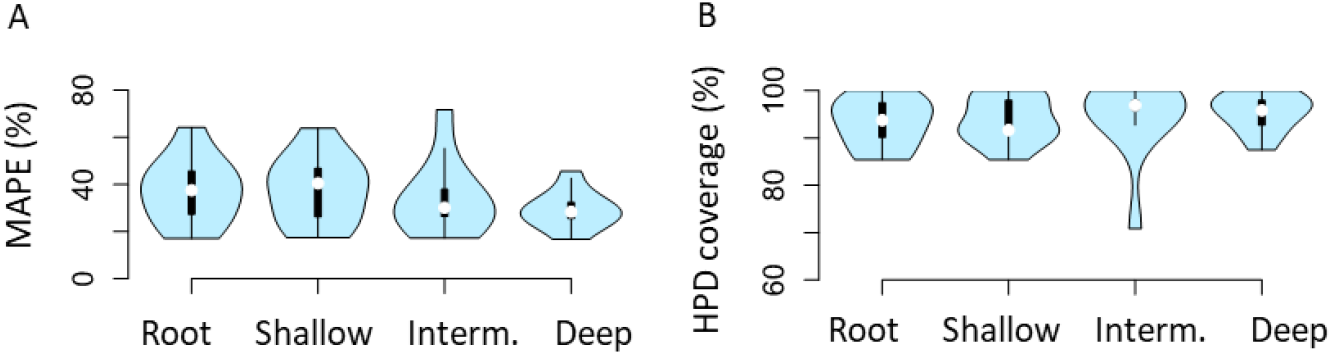
Usefulness of adding one calibration to ddBD tree prior analysis. Distribution of (A) mean absolute percent error, MAPE, and (B) HPD coverage for ddBD tree prior estimates across all simulated datasets when one shallow, intermediate (interm.), and deep calibration is used.

### 3.4. Using ddBD tree prior with diffused calibrations

In all the above analyses, calibrations were well-constrained and as used in the original study. However, the fossil-record usually provides excellent minimum bounds on species divergences, but not the maximum bounds (Bromham, 2019; Marshall, 2008; Hedges *et al*., 2018; Parham *et al*., 2012; Warnock *et al*., 2012). One way to avoid this fossilization of molecular derived dates is to use fossil-based node age estimates as minimum-only calibration priors (with diffused uncertainty densities, as appropriate). However, this would have the side effect of making the phylogeny less calibration-rich. Therefore, we examined if the ddBD tree prior is better than flat priors when only diffused (uniform density) calibrations are used.

Similar to the analysis of well-constrained calibrations, the use of only one diffused internal calibration improved node age estimates to be more concordant with the reported times obtained using all 35 calibrations. The average node age difference was 34.4% for ddBD estimates (**Fig. 11A**), and estimated HPDs contained 56.3% of the reported times (**Fig. 11B**). The inclusion of more diffused calibrations changed the results marginally. As expected, the improvement of estimates was smaller than using well-constrained calibrations because diffused calibrations offer less information to calibrate molecular clocks. Performance of flat tree prior was worse than ddBD tree prior when only a few diffused calibrations were used (**Fig. 11**, red vs. blue). We also conducted an analysis using seven random internal calibrations and found that the difference between estimates and the reported values became much smaller for both ddBD and flat tree priors (average error ∼8.4%, HPD coverage ∼96.7%). These results suggest that ddBD tree prior works well with diffused calibrations and can provide good estimates in calibration-rich (seven diffused calibrations) cases. However, the benefit of using ddBD tree prior is more prominent for calibration-poor phylogenies (e.g., 0-4 calibrations).

**Figure 11.**
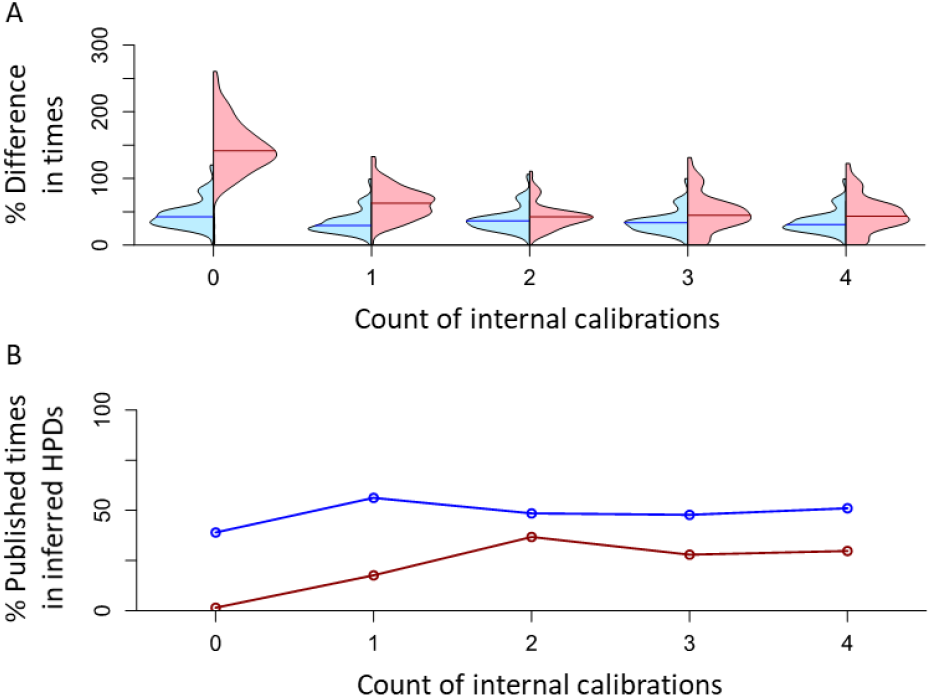
Analysis of empirical data using different numbers of diffused internal calibrations. (A) Distribution of absolute percent differences between published times and times estimated using ddBD (blue) and the flat (red) tree prior when different numbers of internal calibrations are used. Lines in violin plots are median values. (B) Percent nodes whose published times within HPDs obtained using ddBD (blue) and the flat (red) tree prior when different numbers of internal calibrations are used. Zero on x-axis represents the case where only the root calibration and no internal calibrations are used.

### 3.5. Impact of substitution model violation and phylogenetic error on ddBD tree prior

In all the above analyses of simulated datasets, we used the correct substitution model and the correct tree topology to avoid any potential confounding effect. However, trees with phylogenetic errors may be used in estimating the ddBD tree prior because phylogenetic trees inferred from molecular sequences frequently contain topological errors. We found that the accuracy of node times estimated (MAPE and MPE) using ddBD tree priors derived from inferred phylogenies were less than 1% different from those based on the correct phylogenies for every pair of conditions when using the ddBD tree priors (**Fig. 12**), even though inferred phylogenies differed as much as 12.8% from the true tree. While a small amount of phylogeny errors had a limited adverse impact on the usefulness of the ddBD tree prior, the use of an oversimplified model reduced the accuracy of time estimates substantially (as much as 10%), especially when the phylogeny used was incorrect. Importantly, however, overall patterns in **figure 12** make it clear that the main determinant of the accuracy of time estimates was the use of ddBD tree prior, as it resulted in much better estimates than the use of flat tree prior under every condition tested.

**Figure 12.**
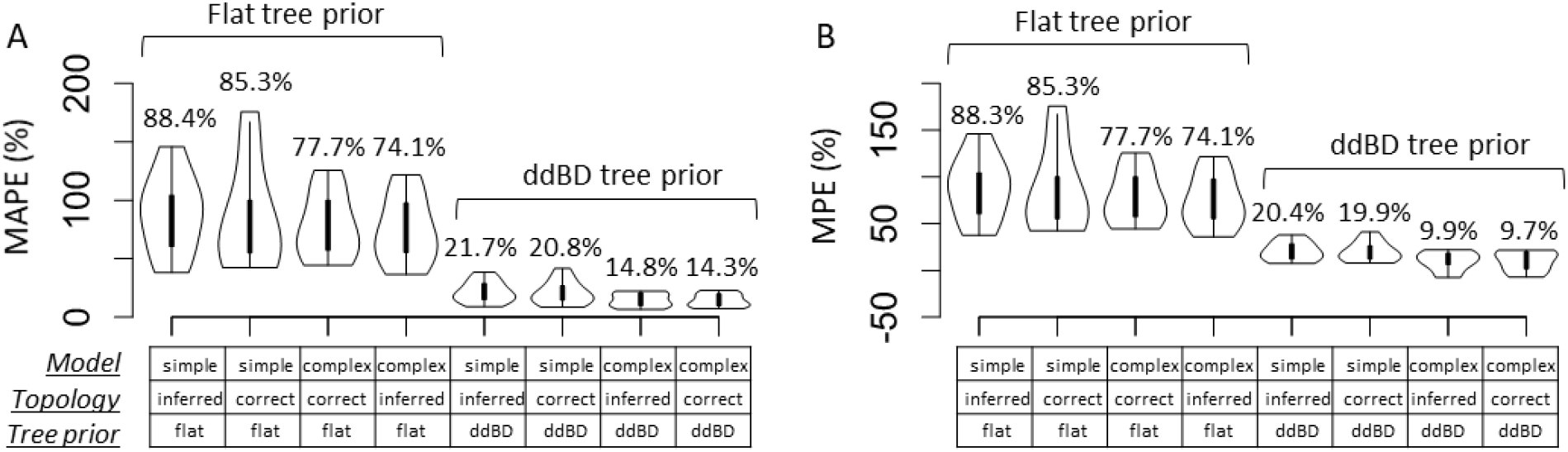
(A) Mean absolute percent error, MAPE, and (B) mean percent error, MPE, of time estimates obtained using different combinations of substitution models (complex or simple), topologies (correct or inferred), and tree priors (ddBD or flat). MAPE and MPE distributions are shown by violin plots. Bars inside violin plots represent the interquartile range. Average values of MAPE and MPE of each violin plot are shown on the top.

## 4. Discussion

Our analyses suggest that using a tree prior inferred from the molecular phylogeny, along with a few well-constrained internal calibrations, can produce time estimates of a quality similar quality to that obtained by using a large number of well-constrained calibrations. Even using a few diffused calibration time priors, which are much easier to derive from the fossil record, coupled with a ddBD tree prior, may produce better estimates than those obtained using the default flat tree prior. These findings indicate that a tree prior derived from data itself can constrain uncertainty densities of diffused calibrations. This result may be explained by the fact that Bayesian time estimates are a function of sequence divergences, clock calibrations, tree prior, and their interactions. The use of an informative prior in Bayesian methods is expected to produce better posterior estimates of node ages and credibility intervals. These findings highlight the advantage of using a ddBD tree prior to dating species divergence times for calibration-poor clades.

Our findings may address a growing concern that the molecular dating using many calibration priors with narrow densities suppress the time structure present in the molecular dataset, causing molecular-derived node ages to become spuriously concordant with the fossil record (Bromham *et al*., 2018; Battistuzzi *et al*., 2015; Hedges *et al*., 2018). One way to avoid this fossilization of molecular derived dates is to use fossil-based node age estimates as minimum-only calibration priors (with diffused uncertainty densities, if desired) and only use a few well-constrained, well-established calibrations. Such a calibration design coupled with ddBD speciation prior is likely to yield good time estimates, based on our results, and avoid the problem of “fossilization” of molecular divergence time estimates.

The use of ddBD tree priors may also enhance the detection of problematic/influential calibrations because methods for such purposes often involve estimating node ages with very few or without any calibrations. For example, Near and Sanderson (2004) estimate node ages with one fixed calibration at a time and calculate the distance between estimated node ages with fossil ages to identify inconsistent calibrations. Battistuzzi *et al*. (2015) developed another protocol that utilizes the linear relationship between node ages estimated with and without internal calibrations to detect the influential calibrations. Therefore, the accuracy of influential calibration detection highly relies on the reliability of node times estimated with poor calibrations. Since the ddBD tree prior performed better than the default flat tree prior for calibration-poor phylogenies in our simulated and empirical phylogenies, it may enable better detection of outliers and inconsistent calibrations.

Although analyses show that our method works well, many sources can potentially bias the results. The ddBD priors, phylogenetic trees, and the best-fit substitution models are all inferred from the same multispecies alignment. Fortunately, ddBD can still improve time estimates even when there are phylogeny errors and the substitution model used is not appropriate. More extensive simulations are needed to confirm these early trends, but they are encouraging nonetheless. In the meantime, one may wish to choose a relatively complex model in their analysis and embrace phylogenetic uncertainty by estimating the tree prior for many phylogenies and using it individually for phylogeny-specific dating use. Results from all phylogenies can then be aggregated in the desired way to generate the final timetree.

It is also important to note that our method does not guarantee that one can recover the exact values for birth rate λ, death rate μ, and species fraction ρ, because many combinations of parameter values can result in the same kernel densities (i.e., distribution of node times) (Louca and Pennell, 2020; Stadler, 2009). The estimated parameters work well because MCMCTree uses the parameters provided as input to generate a distribution of times internally. As long as the three parameters recapitulate the approximate distribution of node times, the tree prior will be informative and well-specified in MCMCTree. Since the RRF approach adopted for generating the distribution of node times relaxes the molecular clock, has a strong theoretical foundation, and demonstrated high accuracy in various simulated datasets (Tamura *et al*., 2018; Tao *et al*., 2020), we expect the tree prior constructed to be effective and informative.

Our current method to estimate ddBD is focused on MCMCTree because it is a fast Bayesian method that works well for large datasets.Since different software packages implement the birth-death model differently, we plan to modify our method to suit other Bayesian dating software (e.g., BEAST (Bouckaert *et al*., 2014)) in the future to provide a direct inference of parameter settings. Nevertheless, we expect the benefits of using a ddBD tree prior may also exist for other Bayesian software. Finally, our analyses did not explore phylogenetic uncertainty because MCMCTree requires a fixed phylogenetic topology in dating analysis.

## 5. Conclusions

We conclude that using an informative tree prior, derived directly from the molecular phylogeny being subjected to relaxed clock analyses, reduces the need to have many well-constrained calibrations to obtain reliable time estimates. It shows improvement in the accuracy and precision of divergence time estimation for calibration-poor datasets. As the paucity of extensive fossil records for most taxonomic groups is a key impediment in building the grand timetree of life, the use of ddBD tree priors would enable the dating of the Tree of Life with greater confidence and higher resolution, which would have important implications for studies of diversification dynamics, phylogeography, and biogeography.

## Acknowledgments

We thank Sudip Sharma for his comments on this manuscript.

